# A new tuna specimen (Genus *Auxis*) from the Duho Formation (Miocene) of South Korea

**DOI:** 10.1101/2024.07.29.605724

**Authors:** Dayun Suh, Su-Hwan Kim, Gi-Soo Nam

## Abstract

A partially preserved caudal vertebrae imprint of a tuna was discovered from the Duho Formation (Miocene) of South Korea. This specimen was assigned to the genus *Auxis* and represents the second record of fossil *Auxis* found in South Korea and in the world. We compared the vertebral morphology of the studied specimen to that of currently known species of *Auxis*, including extinct taxa. However, the specimen could not be assigned to any extant or new species of *Auxis* due to anatomical differences and a lack of comparability. The discovery of a new specimen of *Auxis* aligns with theories of high marine biodiversity in the East Sea (Sea of Japan) and its opening in the Early to Middle Miocene. A widely opened East Sea and upwelling activities might have increased the abundance and diversity of large oceanic fishes such as tunas during the deposition of the Duho Formation. The specimen supports paleoenvironmental interpretations of the Duho Formation as pelagic and subtropical. A taphonomic scenario of the specimen was inferred based on the lack of anal pterygiophores and the leaf imprint on the matrix. The specimen would have been exposed for at least a month in a low-energy sedimentary environment at the deep-sea bottom and would have undergone disintegration before being buried.

## Introduction

The family Scombridae includes mostly epipelagic marine fishes, such as tunas, a large, epipelagic predator (Collette and Nauen, 1983). The four genera of tuna, *Auxis, Euthynnus, Katsuwonus*, and *Thunnus*, form the tribe Thunnini. Among the Thunnini, the genus *Auxis* is an epipelagic, neritic, and oceanic genus found worldwide in tropical and subtropical oceans (Collette and Nauen, 1983). *Auxis* consumes various fishes, crustaceans, cephalopods, and other prey and is preyed upon by large tunas, billfishes, barracudas, sharks, and more (Collette and Nauen, 1983). *Auxis* comprises two extant species: the frigate and bullet tunas (*Auxis thazard* and *Auxis rochei*). They exhibit significant morphological similarities (Vieira et al., 2022) and little osteological differences. The fossil history of *Auxis* is incomplete due to a lack of fossil records and many invalidations of fossil specimens. The fossilization potential is extremely low for taxa in deep-sea and pelagic environments with a 15% and 3% possibility of fossilization respectively due to factors like the fragile skeleton and high environmental stress (Shaw et al., 2020). As a result, *Auxis* fossils are extremely rare, with very few reported specimens. Meanwhile, several fossil specimens previously identified as *Auxis* have undergone multiple taxonomic changes within the Scombridae (Nam et al., 2021). Several cases of *Auxis* misidentification have been noted in the literature. Five scombrid species from the Miocene in Europe were initially identified as *Auxis* by Kramberger-Gorjanović (1882) and Gorjanović-Kramberger (1895); however, Nam et al. (2021) later disputed these identifications based on differences in traits such as vertebral count, lack of the haemal arch, and gill cover and dentition size. Additionally, Woodward (1901) reclassified a scombrid fossil originally described by Agassiz (1833–1844) as *Auxis*, representing what was considered the earliest record of *Auxis* from the Eocene. Bannikov and Sorbini (1984) later corrected this identification, noting discrepancies in vertebral number and shape of haemal arches. Another specimen, a Middle to Upper Miocene scombrid from Russia identified as *Auxis* by Bogatshov (1933), was reclassified by Bannikov (1985) due to an inconsistent vertebral count. With these invalidations, the only currently accepted fossil record of *Auxis* dates back to the Miocene and is reported from the same formation as the specimen described in this paper (†*Auxis koreanus*, Nam et al., 2021). Moreover, the detailed study of the vertebral anatomy of *Auxis* has been hindered by the paucity of recovered specimens including both skulls and vertebrae.

An imprint of tuna vertebrae was collected from the Duho Formation, Pohang City, South Korea, in 2020 (Fig. 1). The new specimen (GNUE322001, Gongju National University of Education) represents the second discovery of *Auxis* from the Duho Formation of the Korean Peninsula and the second valid *Auxis* specimen in the world. Although the specimen is preserved poorly and lacks cranial elements, it possesses diagnostic characters of the vertebrae of the genus *Auxis*: the bifurcated inferior antero-zygapophysis with a long pedicle and no trellis. This paper describes this new tuna specimen and discusses the palaeoecological implications of the presence of tunas in the Miocene of South Korea.

**Figure 1.**
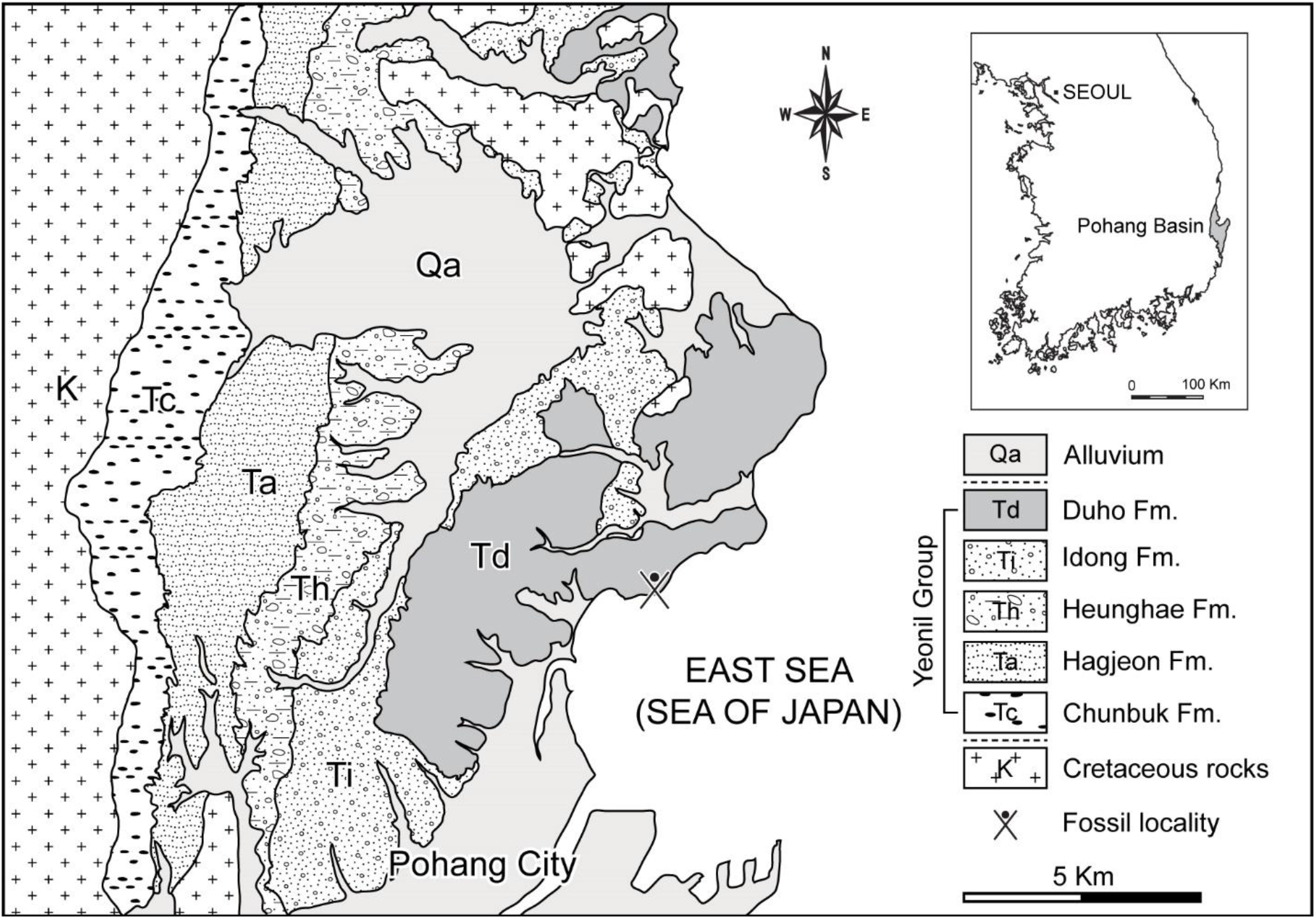
Geologic map of the northern part of the Pohang area with Cenozoic basins in South Korea (inset), depicting the fossil locality where GNUE322001 was collected.

### Geological setting

The Pohang Basin is the largest Cenozoic basin in South Korea (Yoon, 1975; Fig. 1) and is a pull-apart basin that started to form by post-volcanism subsidence at ~17 Ma (Sohn et al., 2001). The Yeonil Group, in the Pohang Basin, includes more than 1 km thick non-marine to deep-marine strata that are characterized predominantly of clastic sediments of marine origin (Sohn et al., 2001; Kim, 2008). This group comprises conglomerates and sandstones along the basin margin and hemipelagic mudstones and sandstones towards the basin center (Sohn et al., 2001; Woo and Khim, 2006). The Yeonil Group consists of the Duho, Idong, Heunghae, Hagjeon, and Chunbuk Formations (Yoon, 1975; Yun, 1986; Fig. 1). The Duho Formation, where the studied specimen was collected, occurs in the uppermost part of the Yeonil Group and is about 250 m thick (Yun, 1986). A pale grey to light brown homogeneous mudstone with intercalated sandstone is the main deposit of the Duho Formation (Hwang et al., 1995; Kim and Paik, 2013). The Duho Formation produces a variety of marine invertebrate and vertebrate fossils, including mollusks (Kim and Lee, 2011; Kong and Lee, 2012), fishes (Ko, 2016; Ko and Nam, 2016; Kim et al., 2018; Nam et al., 2019; Nam et al., 2021; Nam and Nazarkin, 2022; Nazarkin and Nam, 2022; Malyshkina et al., 2023; tab. 1), and whales (Lim, 2005; Lee et al., 2012). Such a diverse fossil record has produced equally diverse paleoenvironmental interpretations of the depositional environment of the Duho Formation. The paleoenvironmental interpretation of the Duho Formation ranges between shallow marine (Kim, 1965; Yun, 1985), offshore (Lee, 1992; Yoon, 1975; Yoon, 1976), low energy (Seong et al., 2009; Kim and Lee, 2011), hemipelagic (Chough et al., 1990; Kim and Paik, 2013), and deep-sea environments (Chough et al., 1990; Kim and Paik, 2013). Various studies on the age of the Duho Formation additionally resulted in diverse interpretations (Kim et al., 2018), ranging from the Early Miocene based on Zircon dating (Lee et al., 2014), Middle Miocene based on paleomagnetic dating and volcanic rocks (Kim et al., 1993; Chung and Koh, 2005), and Late Miocene based on dinoflagellate and radiolarian fossils (Byun and Yun, 1992; Bak et al., 1996).

**Table 1.** List of fish taxa from the Duho Formation.

| Taxa |  | References |
| --- | --- | --- |
| Actinopterygii | Pleuronectiformes | Ko (2016) |
|  | *Pleuronichthys* sp. | Ko and Nam (2016) |
|  | †*Vinciguerria orientalis* | Nam et al. (2019) |
|  | †*Stenobranchius sangsunii* | Nam and Nazarkin (2022) |
|  | *Vinciguerria* sp. | Nazarkin and Nam (2022) |
|  | †*Auxis koreanus* | Nam *et al.* (2021) |
| Elasmobranchii | †*Carcharodon hastalis* | Kim *et al.* (2018)<br>Malyshkina *et al.* (2023) |
|  | *Hexanchus griseus* | Malyshkina *et al.* (2023) |
|  | †*Dalatias orientalis* | Malyshkina *et al.* (2023) |
|  | *Mitsukurina owstoni* | Malyshkina *et al.* (2023) |
|  | †*Otodus megalodon* | Malyshkina *et al.* (2023) |
|  | †*Parotodus benedenii* | Malyshkina *et al.* (2023) |
|  | †‘*Isurus*’ *planus* | Malyshkina *et al.* (2023) |
|  | *Isurus* sp. | Malyshkina *et al.* (2023) |
|  | †*Cetorhinus huddlestoni*, | Malyshkina *et al.* (2023) |
|  | *Carcharhinus* aff. *C. plumbeus* | Malyshkina *et al.* (2023) |
|  | *Carcharhinus* aff. *C. amblyrhynchos* | Malyshkina *et al.* (2023) |
|  | *Carcharhinus* aff. *C. altimus* | Malyshkina *et al.* (2023) |
|  | †*Galeocerdo aduncus* | Malyshkina *et al.* (2023) |

## Materials and methods

The specimen GNUE322001, a partially preserved caudal tuna vertebrae imprint, is housed in the Gongju National University of Education (GNUE), Gongju City, South Korea. The specimen was photographed using a digital camera (Sony A7R4A). Image processing and line drawings of the specimen were done using Adobe Photoshop v 23.4.2. and Adobe Illustrator v 26.4.1. All measurements were taken using a digital caliper.

### Anatomical nomenclature

We follow the terminology of Starks (1910), which was applied to *Auxis*, to describe peculiar vertebral structures of the studied specimen and occasionally refer to the terminology of Romeo and Mansueti (1962) for efficient comparison between *Auxis, Euthynnus*, and *Katsuwonus* of the tribe Thunnini.

### Repositories and institutional abbreviation

The specimen is deposited in the Gongju National University of Education (GNUE), Gongju City, South Korea.

## Results

### Systematic Paleontology

Order Perciformes Nelson, 2006

Suborder Scombroidei Nelson, 2006

Family Scombridae Rafinesque, 1815

Tribe Thunnini Starks, 1910

Genus *Auxis* Cuvier, 1829

### Type species

*Scomber rochei* Risso, 1810

### Occurrence

Duho Formation, Hwanho-dong, Buk-gu, Pohang City, North Gyeongsang Province, South Korea (N36°3’49.10”, E129°23’47.07”) (Fig. 1), preserved in a massive grey mudstone in the Duho Formation (Fig. 2).

**Figure 2.**
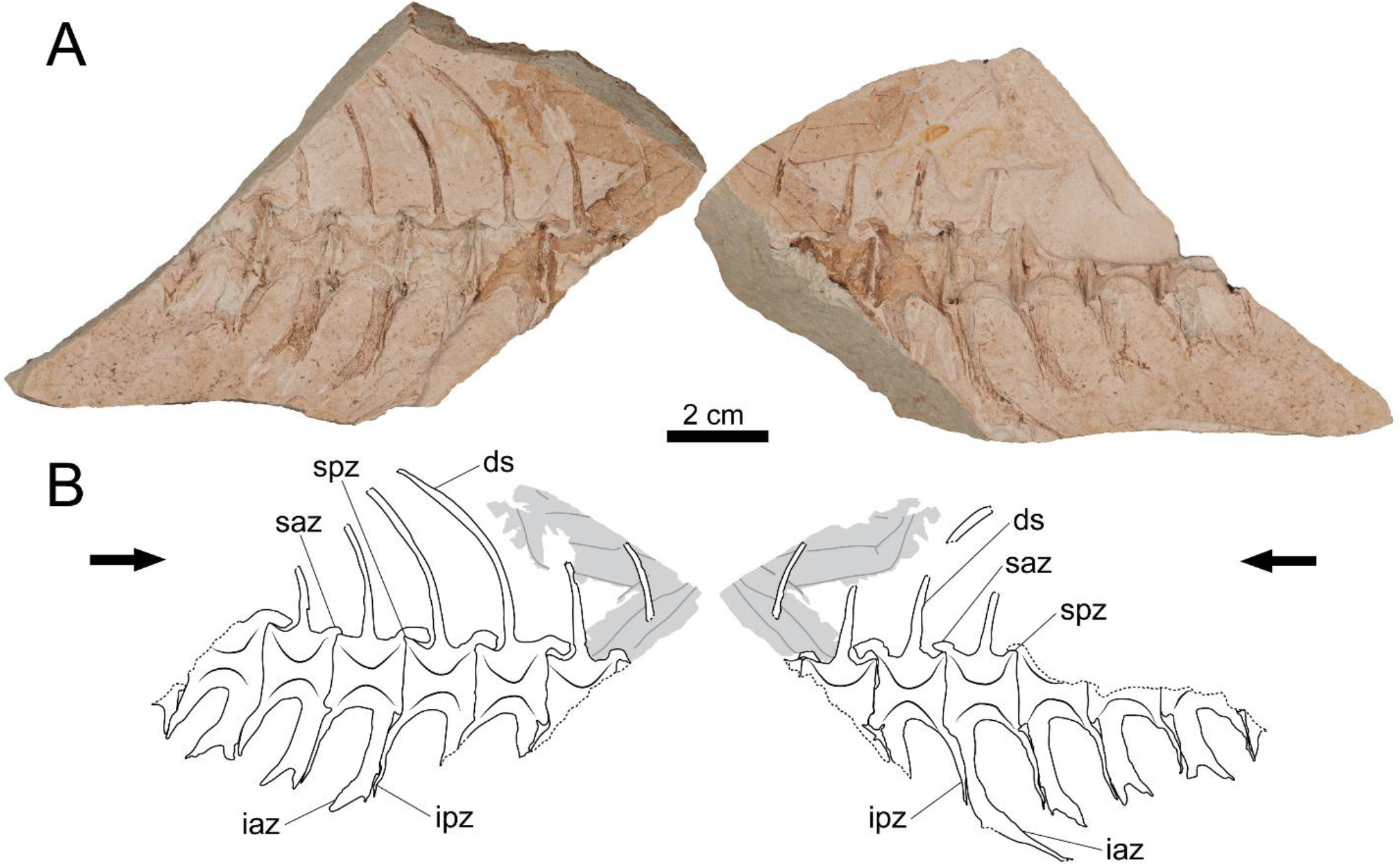
Photographs and drawings of GNUE322001. (A) Photographs of GNUE322001. Each counterpart mold shows a lateral side of the vertebrae without the original bones. (B) Drawings of GNUE322001. Black arrows point towards the anterior direction of the vertebrae. Dashed lines indicate a broken edge. Grey areas indicate an unidentified leaf imprint. Vertebral terminology follows Starks (1910). Abbreviations: ds, dorsal spine; iaz, inferior antero-zygapophysis; ipz, inferior postero-zygapophysis; saz, superior antero-zygapophysis; spz, superior postero-zygapophysis.

### Description

Due to the dissolution of the original bones, only the molds of the eight articulated caudal vertebrae are partially preserved (Fig. 2). In particular, due to the breakage of the matrix, only small fragments of the first and last vertebrae are preserved. The centra have an amphicoelous shape, consisting of two robust cones, and both cones are connected by what appears as a wide foramen. In all Thunnini and certain fish vertebrae, the centrum is not pierced thus lacks a notochordal foramen (Starks, 1910; Graham and Dickson, 2000). Therefore, the apparent wide foramen in GNUE322001 is interpreted as the result of the split of the specimen along a parasagittal plane, and does not represent a real anatomical trait. The anteroposterior length and dorsoventral height of the centrum are subequal, and the dorsal and ventral margins of the centrum are slightly concave in lateral view.

The superior antero-zygapophysis is quite large and dorsoventrally deep, covering most of the posterodorsal margin of the preceding centrum from the posterior margin of the centrum to the posterior edge of the base of its dorsal spine (Fig. 2). In contrast, the superior postero-zygapophysis is weakly developed and is barely discernable in lateral view due to the overlapping superior antero-zygapophysis of the following vertebra.

The dorsal spine originates from the centrum at mid-length, and is slightly angled posteriorly, forming an angle of ~80-85° with the posterodorsal margin of the centrum (Fig. 2). It slightly curves posteriorly at a third of the total length of the preserved spine from its base.

On the fourth to seventh vertebrae, the preserved inferior antero- and postero-zygapophyses project from the centrum ventroposteriorly at an angle of ~70-80° (Fig. 2). The length of these ventral processes of the vertebrae progressively decreases in more posterior vertebral positions. The length of these processes in the first to third vertebrae cannot be assessed due to incomplete preservation.

All preserved inferior antero-zygapophyses are bifurcated into anterior and posterior branches, and the latter tends to be longer (Fig. 2). The inferior antero-zygapophysis of the fourth vertebra is much longer than that of the other vertebrae. It extends nearly to the level of the posterior tip of that of the following vertebra. The inferior postero-zygapophysis extends almost to the ventral tip of the anterior branch of the inferior antero-zygapophysis of the following vertebra. They firmly attach to each other along the entire posterior margin of the inferior postero-zygapophysis.

### Remarks

We classify GNUE322001 as anterior to the middle caudal vertebrae based on the progressively decreasing lengths of the ventral processes, a pattern also observed in the caudal vertebrae of extant *Auxis* (see Uchida, 1981: fig. 24; Jawad et al., 2013: fig. 1; Fig. 4).

The classification of extant *Auxis* is based primarily on the relative body depth, corset width, the number of gill rankers and color pattern (Collette and Aadland, 1996). The extinct *Auxis*, †*A. koreanus*, is distinguished from extant *Auxis* by the osteological differences in skull elements (Nam et al., 2021). Because only the caudal vertebrae are preserved in GNUE322001, the skull is not available for comparison with other species of *Auxis*. However, GNUE322001 exhibits several distinct morphological features in the caudal vertebrae, which we compare with that of other extant *Auxis* species in the following discussion.

## Discussion

### Anatomical comparisons

Among the Thunnini, the genera *Auxis, Euthynnus*, and *Katsuwonus* share a morphological similarity in the inferior antero-zygapophysis in that it is bifurcated into anterior and posterior branches, a unique characteristic only observed in these three genera. However, *Auxis* exhibits ventral bifurcation only in the caudal vertebrae, whereas this trait is also found in the posterior abdominal vertebrae in *Euthynnus* and *Katsuwonus* (see Godsil and Byers 1944: fig. 19; Godsil, 1954: fig. 83; Yoshida and Nakamura, 1965: fig. 3). Furthermore, the pedicle of *Auxis*, a median rod formed by the fusion of both sides of the inferior antero-zygapophyses below the centrum and above the haemal canal (Kishinouye, 1923), is far longer than that of *Euthynnus* and *Katsuwonus* (Godsil, 1954; Fig. 3).

**Figure 3.**
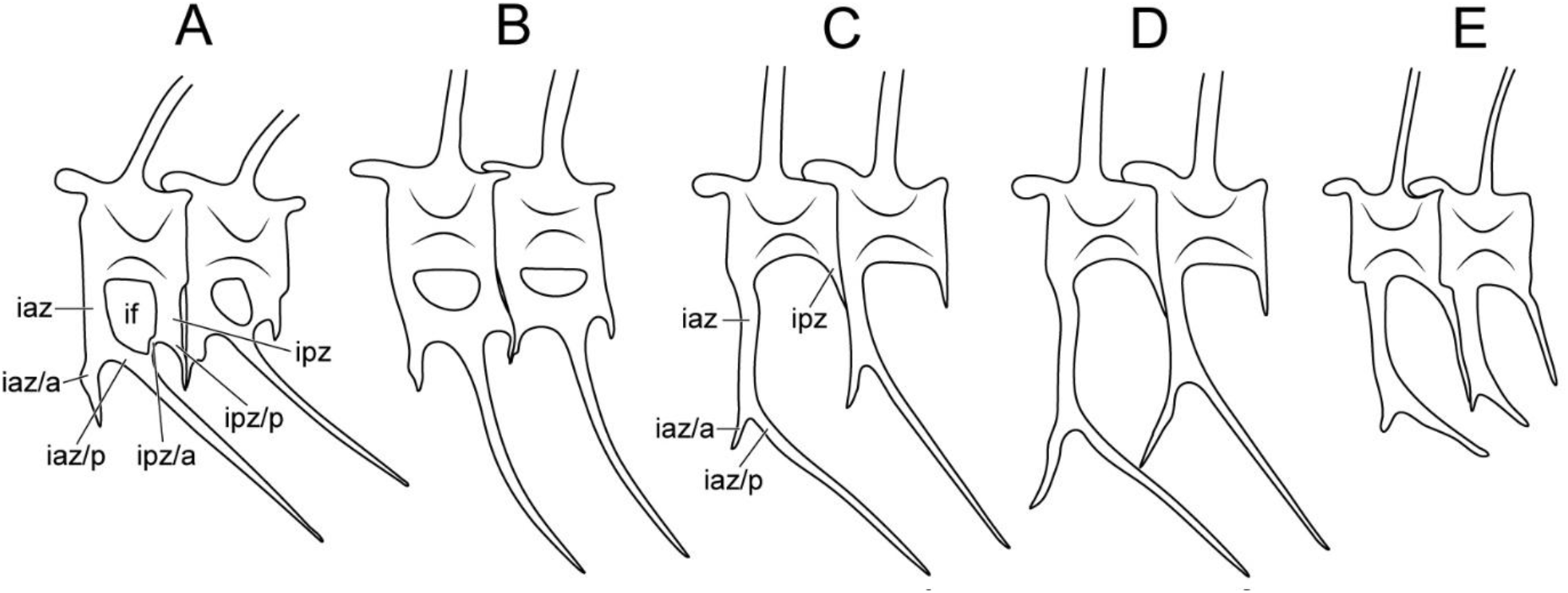
Comparative diagram of the anterior caudal vertebrae of *Auxis, Euthynnus, Katsuwonus*, and GNUE322001. (A) *Euthynnus*. (B) *Katsuwonus*. (C) *A. rochei*. (D) *A. thazard*. (E) GNUE322001 (Godsil and Byers, 1944; Yoshida, 1979; Uchida, 1981). Vertebral terminology follows Starks (1910). Abbreviations: iaz/a, anterior branch of inferior antero-zygapophysis; iaz/p, posterior branch of inferior antero-zygapophysis; ipz/a, anterior branch of inferior postero-zygapophysis; ipz/p, posterior branch of inferior postero-zygapophysis; if, inferior foramen. Note that in *Euthynnus* (A) and *Katsuwonus* (B), the posterior branch of inferior antero-zygapophysis and the anterior branch of inferior postero-zygapophysis are fused, forming the inferior foramen and trellis. Size not to scale.

**Figure 4.**
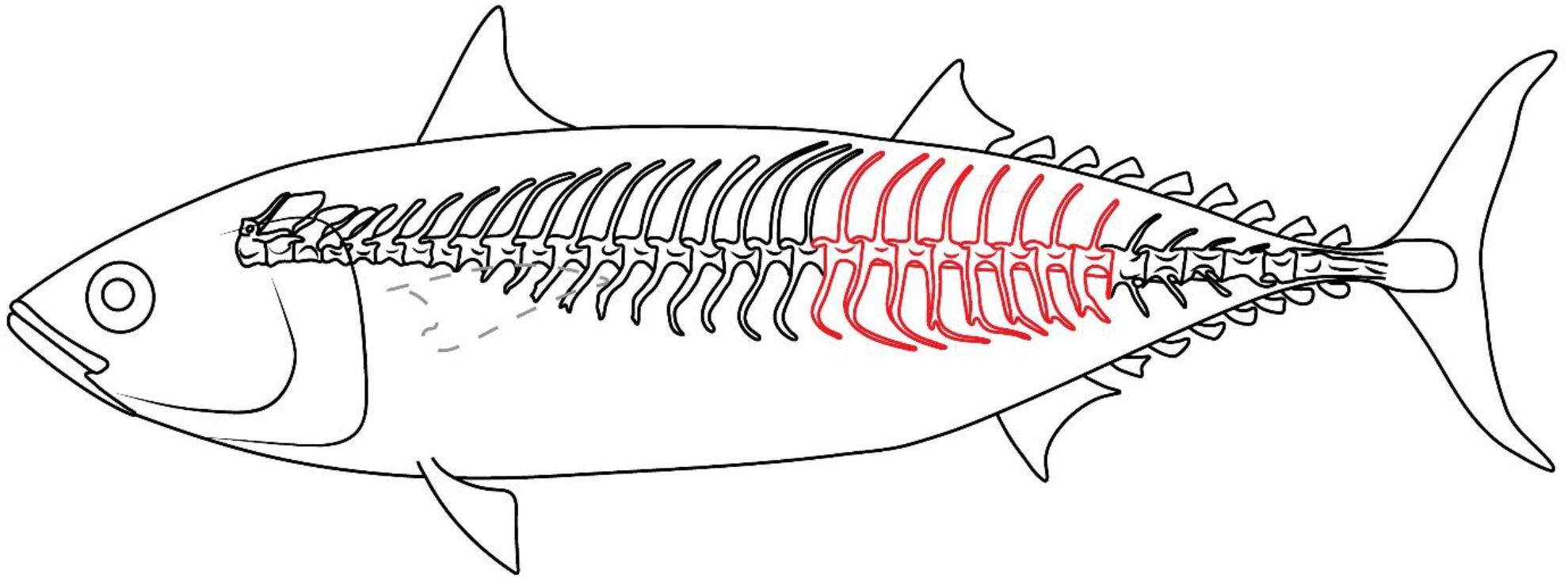
Reconstruction of GNUE322001. The red-lined vertebrae indicate the possible position of the vertebrae of GNUE322001 in the vertebral column.

Most significantly, the anterior caudal vertebrae in *Euthynnus* and *Katsuwonus* are characterized by the presence of an inferior foramen and trellis pattern. The inferior foramen is a hole-like structure fenestra created by the complete fusion of the posterior branch of the inferior antero-zygapophysis (prehaemapophysis of Romeo and Mansueti, 1962) and the anterior branch of the inferior postero-zygapophysis (posthaemapophysis of Romeo and Mansueti, 1962) under the centrum (see Romeo and Mansueti, 1962: fig. 2D; Fig. 3A, B). The trellis pattern is formed by the continuous repetition of the inferior foramen across the vertebrae (Fig. 3A, B). In *Auxis*, the inferior foramen and trellis pattern are scarcely developed, and even when present, only occur in the posterior caudal vertebrae, unlike in *Euthynnus* and *Katsuwonus* (Kishinouye, 1923; Godsill, 1954; Fig. 3).

Although the cranial elements are not preserved in GNUE322001, this specimen was identified as *Auxis* primarily based on having a bifurcated inferior antero-zygapophysis with a long pedicle and no trellis. Based on the vertebral column of extant *Auxis* (see Uchida, 1981: fig. 24; Jawad et al., 2013: fig. 1), it is suggested that GNUE322001 represents the anterior to the middle caudal vertebral series (Fig. 4) as indicated by the lengths of ventral processes that progressively decrease throughout the vertebral series in this taxon.

There are three valid taxa within *Auxis*, including an extinct species (*A. thazard, A. rochei*, and †*A. koreanus*) (Collette and Aadland, 1996; Nam et al., 2021). GNUE322001 is morphologically similar to the vertebrae of *A. rochei* in that the anterior branch of the inferior antero-zygapophysis is short and does not reach the preceding inferior antero-zygapophysis (Yoshida and Nakamura, 1965; Uchida, 1981; Fig. 3C, E). In *A. thazard*, the anterior branches of the inferior antero-zygapophyses are long enough to contact the preceding inferior antero-zygapophyses (Fig. 3D). Comparisons with the extinct taxon †*A. koreanus* are not possible because the caudal vertebrae are unknown in this species, as they are not preserved in its only known specimens (Nam et al., 2021). Although †*A. koreanus* was also discovered in the Duho Formation, as was GNUE322001, it is challenging to assign GNUE322001 to †*A. koreanus* based solely on their shared occurrence within the same formation. Furthermore, the centra of †*A. koreanus* and GNUE322001 differ significantly in size, with lengths of approximately 0.5 and 1.5 cm respectively (Nam et al., 2021; Fig. 2). Therefore, additional study and discovery of *Auxis* specimens from the Duho Formation are necessary to determine the relationship between GNUE322001 and †*A. koreanus*, as well as other extant species of *Auxis*.

### Paleoenvironmental perspectives

The major opening of the East Sea that occurred between 23 and 18 Ma widened the gap between the Japanese Arc and the Korean Peninsula by 200-250 km (Sohn et al., 2001). The opening of the East Sea would have facilitated the creation of a variety of marine environments. This increased environmental complexity likely provided habitats that were able to support a wider range of marine species. A diverse fossil record of large oceanic animals in the East Sea during this period such as the tunas (Nam et al., 2021; GNUE322001 in this paper), sharks (Kim et al., 2018), and whales (Lim, 2005; Lee et al., 2012) aligns with this theory.

Upwelling regions, although only constituting 0.1% of the total ocean areas (Wang and Lee, 2019), are where fishes are most abundant due to high production rates (Lalli and Parsons, 1997). One of such fishes is the tuna, which are attracted by the zones of foraging availability created by upwelling zones (Grandperrin, 1978; Nicol et al., 2014). Additionally, based on the record of the fossilized diatom resting spores, which indicate an upwelling activity in the Duho Formation (Hargraves, 1979; Lee, 1993), Kim et al. (2018) hypothesized that the biodiversity of the East Sea increased due to the influence of upwelling during the deposition of the Duho Formation. Thus, it can be concluded that upwelling activity during the Miocene increased pelagic fishes’ and their preys’ biodiversity in the East Sea.

Today, tunas inhabit tropical and subtropical epipelagic ocean ecosystems (Collette and Nauen, 1983). During warm seasons, tunas move into coastal areas, whereas in colder seasons, they exclusively occupy deeper offshore waters (Kishinouye, 1923). In deep epipelagic environments, tunas specifically inhabit deep rocky banks and forage in the deep scattering layer (Kishinouye, 1923; Graham and Dickson, 2004). Graham and Dickson (2004) argue that most oceanic physical and biological features observed in ocean ecosystems today have been in place since the Miocene. This suggests that the East Sea during the Miocene likely supported a pelagic and subtropical environment, as tuna, which inhabit such ecosystems, were present.

This interpretation is supported by researchers who consider the Duho Formation to represent a deep-sea and subtropical environment, among the various interpretations of its environmental context (see the Geological Setting section). Recent discoveries of pelagic sharks (Malyshkina et al., 2023), lightfishes (Nam et al., 2019; Nazarkin and Nam, 2022), and lanternfish (Nam and Nazarkin, 2022) provide evidence that the Duho Formation represents a deep-sea environment. Moreover, based on discoveries of certain plant fossils, some researchers interpret the Duho Formation to have been a subtropical environment (Kim, 2008, 2010; Kim et al., 2009, 2017; Jung and Lee, 2009). The discovery of GNUE322001 supports both of these interpretations.

### Taphonomic interpretations

The absence of anal pterygiophores in GNUE322001, which in tunas are located directly under the prehaemapophyses (Fig. 2), suggests that the specimen underwent significant decomposition underwater. The first steps of decomposition of a fish involve the disarticulation of the jaw and external scales as soft tissues (muscles, skins) decompose (Burrow and Turner, 2012). However, body parts are often disarticulated but still loosely connected (Burrow and Turner, 2012). At this stage, invertebrate and vertebrate scavengers completely disconnect the bones by feeding on the soft tissue or the bones themselves (Burrow and Turner, 2012). In GNUE322001, the absent anal pterygiophores would have been disconnected and/or consumed by marine scavengers, indicating that the vertebrae would have been underwater for a long time. However, the exact taphonomic time frame cannot be determined with the partially preserved vertebrae.

An unidentified leaf imprint is preserved on the anterior portion of the vertebrae of GNUE322001 (Fig. 2). Since the fine-grained matrix indicates that the specimen was buried in a low-energy sedimentary environment at the deep-sea bottom, the leaf associated with GNUE322001 would have traveled from shore to the depths of the sea. The leaf exhibits tears on its edges, a characteristic of the fragmentation stage of decomposition where marine detritivorous invertebrates feed on deposited leaves (Bridgham and Lamberti, 2009). The decomposition rate during fragmentation varies depending on salinity; aquatic ecosystems with lower salinity are correlated with faster decomposition (Quintino et al., 2009). Thus, decay rates are greatest in freshwater ecosystems, followed by transitional communities, and slowest in marine ecosystems (Quintino et al., 2009). While the torn edges of the leaf imprint associated with GNUE322001 resemble those of leaves that underwent a two-week decomposition in transitional communities (Bridgham and Lamberti, 2009: fig. 15.2), leaves deposited in marine ecosystems take more than twice the time to exhibit a similar amount of biomass remain (Quintino et al., 2009: fig. 4). Thus the leaf associated with GNUE322001 would have decomposed after a month of being exposed to water. Although the vertebrae and leaf have experienced different decompositions in isolated conditions, based on the taphonomic time frame inferred from the preservation of the leaf imprint, it can be estimated that the decomposition of GNUE322001 took at least a month. However, perfectly preserved leaves were also reported from the Duho Formation (Jung and Lee, 2009); therefore, the taphonomic scenario inferred from GNUE322001 does not represent a general depositional condition of the Duho Formation.

## Acknowledgments

We thank Professor Yuong-Nam Lee for improving the quality of this manuscript. We also thank two anonymous reviewers for their valuable feedback and Dr. Adriana López-Arbarello for overseeing the review process and providing valuable suggestions.

## Funding

This work was not supported by any funding source.

## Conflict of interest disclosure

The authors declare that they comply with the PCI rule of having no financial conflicts of interest in relation to the content of the article.

## Author contributions

Dayun Suh contributed to conceptualization, formal analysis, investigation, visualization, writing of the original draft, and writing of review and editing. Su-Hwan Kim contributed to conceptualization, formal analysis, investigation, methodology, supervision, validation, visualization, and writing of review and editing. Gi-Soo Nam contributed to resources, and validation, and writing of review and editing.

## Notes

### Competing Interest Statement

The authors have declared no competing interest.

### Summary of Updates

The final version of the manuscript, including a typeset PCI Paleo cover page.

## References

Agassiz, L., 1833-1844, Recherches sur les poissons fossiles, 5 Volumes: Neuchâtel, Imprimerie de Petitpierre, 1420 p.

Bak, Y., Lee, J.D., and Yun, H., 1996, Middle Miocene radiolarians from the Duho Formation in the Pohang Basin, Korea: Journal of the Paleontological Society of Korea, v. 12, p. 225– 261.

Bannikov, A.F., 1985, Iskopayemye skumbriyevye SSSR. Trudy Paleontologicheskogo instituta AN SSSR, v. 210, p, 1–111 [In Russian].

Bannikov, A.F. and Sorbini, L., 1984, On the occurrence of genus Scombrosarda in the Eocene of Bolca, v. 4, p. 307–317.

Bogatshov, V.V., 1933, Materialy po izucheniyu tretichnoi ikhtiofauny Kavkaza. Trudy Azerbaidzhanskogo Neftyanogo Issledovatelskogo Instituta, Geologicheskii Otdel, v. 15, p. 1–62 [In Russian].

Bridgham, S.D., and Lamberti, G.A., 2009, Ecological dynamics III: Decomposition in wetlands, in Maltby, E., and Barker, T., eds., The Wetlands Handbook: Blackwell Publishing, Oxford, p. 326–345. 10.1002/9781444315813.ch15

Burrow, C.J., and Turner, S., 2012, Fossil fish taphonomy and the contribution of microfossils in documenting Devonian vertebrate history, in Talent, J.A., ed., Earth and Life: Springer, New York, p. 184–223. 10.1007/978-90-481-3428-1_8

Byun, H., and Yun, H., 1992, Miocene dinoflagellate cysts from the central part of the Pohang Basin, Korea: Journal of the Paleontological Society of Korea, v. 8, p. 164–235.

Chough, S.K., Hwang, I.G., and Choe, M.Y., 1990, The Miocene Doumsan fan-delta, southeast Korea: a composite fan-delta system in back-arc margin: Journal of Sedimentary Research, v. 60, p. 445–455. 10.1306/212f91ba-2b24-11d7-8648000102c1865d

Chung, C.-H., and Koh, Y.-K., 2005, Palynostratigraphic and palaeoclimatic investigations on the Miocene deposits in the Pohang area, South Korea: Review of Palaeobotany and Palynology, v. 135, p. 1–11. 10.1016/j.revpalbo.2005.02.002

Collette, B.B., and Nauen, C.E., 1983, Scombrids of the world, an annotated and illustrated catalogue of tunas, mackerels, bonitos, and related species known to date: Food and Agriculture Organization of the United Nations, Rome, v. 2, 137 p.

Collette, B.B., and Aadland, C.R., 1996, Revision of the frigate tunas (Scombridae, Auxis), with descriptions of two new subspecies from the eastern Pacific. Fishery Bulletin: v. 94, p. 423–441.

Cuvier, G., 1829, Le Règne Animal, distribué d’après son organisation, pour servie de base à l’histoire naturelle des animaux et d’introduction à l’anatomie comparé: Paris, 532 p.

Godsil, H.C., and Byers, R.D., 1944, A systematic study of the Pacific tunas: California Department of Fish and Game, Fish Bulletin, v. 60, p. 1–131.

Godsil, H.C., 1954, A descriptive study of certain tuna-like fishes: California Department of Fish and Game, Fish Bulletin, v. 97, p. 1–185.

Gorjanović-Kramberger, D., 1895, Fosilne ribe Komena, Mrzleka, Hvara i M. Libanona i dodatak o oligocenskim ribama Tüffera, Zagora i Trifalja. Djela Jugoslavenske Akademije Znanosti i Umjetnosti, v. 16, p. 1–67.

Graham, J. B., and Dickson, K. A., 2000, The evolution of thunniform locomotion and heat conservation in scombrid fishes: new insights based on the morphology of Allothunnus fallai: Zoological Journal of the Linnean Society, v. 129, p. 419–466. 10.1111/j.1096-3642.2000.tb00612.x

Grandperrin R., 1978, Influence of currents on the production of tropical seas: consequences for fisheries: SPC Fisheries Newsletter, v. 17, p. 14–20.

Hargraves, P.E., 1979, Studies on marine plankton diatoms, IV. Morphology of Chatoceros resting spores: Beihefte zur Nova Hedwigia, v. 64, p. 99–120.

Hwang, I.G., Chough, S.K., Hong, S.W., and Choe, M.Y., 1995, Controls and evolution of fan delta systems in the Miocene Pohang Basin, SE Korea: Sedimentary Geology, v. 98, p. 147– 179.

Jawad, L.A., Al-Hassani, L., and Al-Kharusi, L.H., 2013, On the morphology of the vertebral column of the frigate tuna, Auxis thazard (Lacepedea, 1800)(Family: Scombridae) collected from the sea of Oman: Acta Musei Nationalis Pragae, v. 69, p. 101–105. 10.14446/amnp.2013.101

Jung, S.H., and Lee, S.J., 2009, Fossil-Winged Fruits of Fraxinus (Oleaceae) and Liriodendron (Magnoliaceae) from the Duho Formation, Pohang Basin, Korea. Acta Geologica Sinica, v. 83, p. 845–852. 10.1111/j.1755-6724.2009.00113.x

Kim, B.K., 1965, The stratigraphic and paleontologic studies on the Tertiary (Miocene) of the Pohang area, Korea: Seoul University Journal Science and Technology Series, v. 15, p. 32–121.

Kim, D.H., and Lee, S.-J., 2011, Fossil scallops from the Hagjeon Formation and the Duho Formation, Pohang Basin: Journal of the Geological Society of Korea, v. 47, p. 235–244 [in Korean with English abstract].

Kim, J., and Paik, I.S., 2013, Chondrites from the Duho Formation (Miocene) in the Yeonil Group, Pohang Basin, Korea: Occurrences and paleoenvironmental implications: Journal of the Geological Society of Korea, v. 49, p. 407–416 [in Korean with English abstract]. 10.14770/jgsk.2013.49.3.407

Kim, J.H., 2008, A new species of Acer samaras from the Miocene Yeonil Group in the Pohang Basin, Korea: Geosciences Journal, v. 12, p. 331–336. 10.1007/s12303-008-0033-6

Kim, J.H., Lee, S., An, J., Lee, H., and Hong, H., 2009, Albizia Fruit Fossils from the Miocene Duho Formation of Yeonil Group in the Pohang Basin, Korea. Journal of the Korean Earth Science Society, v. 30, p. 10–18 [in Korean with English abstract]. 10.5467/jkess.2009.30.1.010

Kim, J.-H., Nam, K.-S., and Jeon, Y.-S., 2017, Diversity of Miocene fossil Acer from the Pohang Basin, Korea: Journal of the Geological Society of Korea, v. 53, p. 387–405 [in Korean with English abstract].

Kim, K.H., Doh, S.-J., Hwang, C.-S., and Lim, D.S., 1993, Paleomagnetic study of the Yeonil Group in Pohang basin: Journal of Korean Society of Economic and Environmental Geology, v. 26, p. 507–518 [in Korean with English abstract].

Kim, S.-H., Park, J.-Y., and Lee, Y.-N., 2018, A tooth of Cosmopolitodus hastalis (Elasmobranchii: Lamnidae) from the Duho Formation (middle Miocene) of Pohang-si, Gyeongsangbuk-do, South Korea. Journal of the Geological Society of Korea: v. 54, p. 121–131 [in Korean with English abstract]. 10.14770/jgsk.2018.54.2.121

Kishinouye, K., 1923, Contributions to the comparative study of the so-called scombrid fishes: Journal of the College of Agriculture, Imperial University of Tokyo, v. 8, p. 293–475.

Ko, J.Y., 2016, The description of the flat fish (Pleuronectiformes) fossils from the Miocene Duho Formation, Pohang Yeonam-dong in Korea and its implication: Journal of the Korean Earth Science Society, v. 37, p. 1–10 [in Korean with English abstract]. 10.5467/jkess.2016.37.1.1

Ko, J.Y., and Nam, K.-S., 2016, Pleuronichthys sp. fossils (Pleuronectidae) from the Duho Formation, Pohang Uhyeon-dong in Korea: Journal of the Korean Earth Science Society, v. 37, p. 133–142 [in Korean with English abstract]. 10.5467/jkess.2016.37.3.133

Kong, D.Y., and Lee, S.J., 2012, Fossil scaphopods from the Hagjeon Formation and the Duho Formation, the Cenozoic Pohang Basin, Korea: Korean Journal of Cultural Heritage Studies, v. 45, p. 218–231 [in Korean].

Kramberger-Gorjanović, D., 1882, Die jungtertiäre Fischfauna Croatiens. Beitrag zur Paläontologie und Geologie Österreich-Ungarns und des Orients, v. 2, p. 86–136.

Lalli, C. and Parsons, T., 1997, Biological oceanography: an introduction: Oxford, Elsevier Butterworth-Heinemann, 314 p.

Lee, T.-H., Yi, K., Cheong, C.-S., Jeong, Y.-J., Kim, N., and Kim, M.-J., 2014, SHRIMP U-Pb zircon geochronology and geochemistry of drill cores from the Pohang Basin: Journal of the Korean Earth Science Society, v. 23, p. 167–185 [in Korean with English abstract]. 10.7854/jpsk.2014.23.3.167

Lee, Y.G., 1992, Paleontological study of the Tertiary molluscan fauna in Korea: Science Reports of the Institution of Geosciences University of Tsukuba, Section B (geological science), v. 13, p. 15–125.

Lee, Y.G., 1993, The marine diatom genus Chaetoceros Ehrenberg flora and some resting spores of the Neogene Yeonil Group in the Pohang Basin, Korea: Journal of the Paleontological Society of Korea, v. 9, p. 24–52.

Lee, Y.-N., Ichishima, H., and Choi, D.K., 2012, First record of a platanistoid cetacean from the middle Miocene of South Korea: Journal of Vertebrate Paleontology, v. 32, p. 231–234. 10.1080/02724634.2012.626005

Lim, J.-D., 2005, The first dolphin fossil from the Miocene of Korea: Current Science, v. 89, p. 939–940.

Malyshkina, T.P., Ward, D.J., Nazarkin, M.V., Nam, G.-S., Kwon, S.-H., Lee, J.-H., Kim, T.-W., Kim, D.-K., and Baek, D.-S., 2023, Miocene Elasmobranchii from the Duho Formation, South Korea: Historical Biology, v.35, p. 1726–1741. 10.1080/08912963.2022.2110870

Nam, G.-S., Ko, J.-Y., and Nazarkin, M.V., 2019, A new lightfish, †Vinciguerria orientalis, sp. Nov. (Teleostei, Stomiiformes, Phosichthyidae), from the middle Miocene of South Korea: Journal of Vertebrate Paleontology, v. 39, e1625911. 10.1080/02724634.2019.1625911

Nam, G.-S., Nazarkin, M.V., and Bannikov, A.F., 2021, First discovery of the genus Auxis (Actinopterygii: Scombridae) in the Neogene of South Korea: Bollettino della Societa Paleontologica Italiana, v. 60, p. 61–67. 10.4435/BSPI.2021.05

Nam, G.-S., and Nazarkin, M.V., 2022, A new lanternfish (Myctophiformes, Myctophidae) from the Middle Miocene Duho Formation, South Korea: Journal of Vertebrate Paleontology, v. 42, e2121924. 10.1080/02724634.2022.2121924

Nazarkin, M.V., and Nam, G.-S., 2022, Miocene lightfish Vinciguerria sp. (Stomiiformes: Phosichthyidae) from the North-West Pacific with notes on the recent evolution of the genus: Historical Biology, v. 34, p. 102–107. 10.1080/08912963.2021.1900839

Nelson, J.S., 2006, Fishes of the World (fourth edition): Hoboken, John Wiley and Sons, 601 p.

Nicol, S., Menkes, C., Jurado-Molina, J., Lehodey, P., Usu, T., Kumasi, B., Muller, B., Bell, J., Tremblay-Boyer, L., and Briand, K., 2014, Oceanographic characterisation of the Pacific Ocean and the potential impact of climate variability on tuna stocks and tuna fisheries: SPC Fisheries Newsletter, v. 145, p. 37–48.

Quintino, V., Sangiorgio, F., Ricardo, F., Mamede, R., Pires, A., Freitas, R., Rodrigues, A.M., and Basset, A., 2009, In situ experimental study of reed leaf decomposition along a full salinity gradient: Estuarine, Coastal and Shelf Science, v. 85, p. 497–506. 10.1016/j.ecss.2009.09.016

Rafinesque, C.S., 1815, Analyse de la nature, ou tableau de l’univers et des corps organisés: Palermo, 224 p. 10.5962/bhl.title.106607

Risso, A., 1810, Ichthyologie de Nice ou histoire naturelle des poissons du département des Alpes Maritimes: Paris, F. Schoell, 388 p. 10.5962/bhl.title.7052

Romeo, J., and Mansueti, A.J., 1962, Little tuna, Euthynnus alletteratus, in northern Chesapeake Bay, Maryland, with an illustration of its skeleton: Chesapeake Science, v. 3, p. 257–263. 10.2307/1350633

Seong, M.N., Kong, D.-Y., Lee, B.J., and Lee, S.-J., 2009, Cenozoic brittle stars (Ophiuroidea) from the Hagjeon Formation and the Duho Formation, Pohang Basin, Korea: Journal of Korean Society of Economic and Environmental Geology, v. 42, p. 367–376 [in Korean with English abstract].

Shaw, J.O., Briggs, D.E.G., and Hull, P.M., 2020, Fossilization potential of marine assemblages and environments: Geology, v. 49, p. 258–262. 10.1130/g47907.1

Sohn, Y.K., Rhee, C.W., and Shon, H., 2001, Revised stratigraphy and reinterpretation of the Miocene Pohang basinfill, SE Korea: sequence development in response to tectonism and eustasy in a back-arc basin margin: Sedimentary Geology, v. 143, p. 265–285. 10.1016/s0037-0738(01)00100-2

Starks, E.C., 1910, The osteology and mutual relationships of the fishes belonging to the family Scombridae: Journal of Morphology, v. 21, p. 77–99. 10.1002/jmor.1050210103

Uchida, R.N., 1981, Synopsis of biological data on frigate tuna, Auxis thazard, and bullet tuna, A. rochei: Silver Spring, US Department of Commerce, National Oceanic and Atmospheric Administration, National Marine Fisheries Service, 63 p. 10.5962/bhl.title.63109

Vieira, J.M., Costa, P.A., Braga, A.C., São-Clemente, R.R., Ferreira, C.E., and Silva, J.P., 2022, Age, growth and maturity of frigate tuna (Auxis thazard Lacepède, 1800) in the Southeastern Brazilian coast: Aquatic Living Resources, v. 35, p. 11. 10.1051/alr/2022010

Wang, Y.-C., and Lee, M.-A., 2019, Composition and distribution of fish larvae surrounding the upwelling zone in the waters of northeastern Taiwan in summer: Journal of Marine Science and Technology, v. 27, p. 8.

Woo, K.S., and Khim, B.-K., 2006, Stable oxygen and carbon isotopes of carbonate concretions of the Miocene Yeonil Group in the Pohang Basin, Korea: Types of concretions and formation condition: Sedimentary Geology, v. 183, p. 15–30. 10.1016/j.sedgeo.2005.09.005

Woodward, A.S., 1901, Catalogue of the Fossil Fishes in the British Museum (Natural History) Part IV: London, British Museum of Natural History, 636 p. 10.5962/bhl.title.162341

Yoon, S., 1975, Geology and paleontology of the Tertiary Pohang Basin, Pohang District, Korea Part 1, Geology: Journal of the Geological Society of Korea, v. 11, p. 187–214.

Yoon, S., 1976, Geology and paleontology of the Tertiary Pohang Basin, Pohang District, Korea Part 2, Paleontology (Mollusca): Journal of the Geological Society of Korea, v. 12, p. 1– 22.

Yoshida, H.O., and Nakamura, E.L., 1965, Notes on schooling behavior, spawning, and morphology of Hawaiian frigate mackerels, Auxis thazard and Auxis rochei: Copeia, v. 1, p. 111–114. 10.2307/1441254

Yoshida, H.O., 1979, Synopsis of biological data on tunas of the genus Euthynnus: Silver Spring, US Department of Commerce, National Oceanic and Atmospheric Administration, National Marine Fisheries Service, 9 p.

Yun, H., 1985, Some fossil Squillidae (Stomatopoda) from the Pohang Tetiary Basin, Korea: Journal of the Paleontological Society of Korea, v. 1, p. 19–31.

Yun, H., 1986, Emended stratigraphy of the Miocene formations in the Pohang Basin Part I: Journal of the Paleontological Society of Korea, v. 2, p. 54–69.

